# Replication-attenuated r3LCMV vectors potentiate tumor control via IFN-I

**DOI:** 10.1101/2023.12.08.570847

**Authors:** Young Rock Chung, Bakare Awakoaiye, Tanushree Dangi, Slim Fourati, Pablo Penaloza-MacMaster

## Abstract

Viral vectors are being used for the treatment of cancer. Yet their efficacy varies among tumors and their use poses challenges in immunosuppressed patients, underscoring the need for alternatives. We report striking antitumoral effects by a nonlytic viral vector based on attenuated lymphocytic choriomeningitis virus (r3LCMV). We show in multiple tumor models that injection of tumor-bearing mice with this novel vector results in improved tumor control and survival. Importantly, r3LCMV also improved tumor control in immunodeficient Rag1-/- mice. Single cell RNA-Seq analyses, antibody blockade experiments, and KO models revealed a critical role for host IFN-I in the antitumoral efficacy of r3LCMV vectors. Collectively, these data demonstrate potent antitumoral effects by a replication-attenuated LCMV vector and unveil mechanisms underlying its antitumoral efficacy.

## Introduction

Cancer is characterized by immunosuppression, which inhibits the ability of the immune system to clear tumor cells. Immunosuppression is triggered by the recruitment of T regulatory cells, as well as upregulation of inhibitory receptors such as PD-1 and CTLA-4, among other factors. While immune checkpoint therapy can partially revert immunosuppression and result in effective tumor control, only ∼30% of patients respond, highlighting the need for alternative immunotherapies. Viruses have emerged as attractive therapies to overcome immunosuppression during cancer. In particular, oncolytic viruses that preferentially infect and replicate in tumor cells have been explored for cancer immunotherapy (*1*). Currently, an oncolytic virus (talimogene laherparepvec, T-VEC) is approved for melanoma patients (*2*). Although this vector can be effective in some patients with melanoma, adverse effects have been reported following the use of this replicating lytic virus, and not all patients respond. Due to safety concerns, immunocompromised patients are typically excluded from receiving this replicating viral therapy, motivating the development of alternative viral vectors for cancer immunotherapy.

In this study, we explored a non-lytic virus, lymphocytic choriomeningitis virus (LCMV), as a cancer immunotherapy. LCMV can be engineered to serve as a replication-attenuated vector that can deliver foreign antigens to the immune system (*3, 4*). Prior studies have shown that immunization of mice with attenuated LCMV vectors expressing tumor antigens improves tumor control and there is an ongoing trial evaluating the efficacy of attenuated LCMV vectors expressing HPV antigens in patients with HPV16+ metastatic head and neck carcinoma (**#NCT04180215**) (*5–7*). The use of viral vectors expressing a cargo of tumor antigens requires knowledge of specific tumor antigens, which may differ depending on the patient and the type of tumor. In this study we interrogated whether replication-attenuated r3LCMV vectors that do not express any tumor antigen provide antitumor protection. Using multiple tumor models, we show that injection of tumor-bearing mice with r3LCMV vectors results in improved tumor control and prolonged survival.

## Results

### LCMV replication improves tumor control

Due to their high immunogenicity, LCMV vectors have been explored as vaccine candidates for various diseases (*8–10*). In these prior studies, LCMV vectors have been genetically modified to include a foreign antigen to prime antigen-specific immune responses. In our study, we tested whether an LCMV vector that does not express any tumor antigen can confer protection against tumor challenges in mice. We first challenged C57BL/6 mice with 10^6^ B16 melanoma cells and at day 5 post-challenge, we treated tumor-bearing mice intratumorally with 2x10^5^ FFU of a replication-attenuated LCMV vector (r3LCMV) (**Fig. 1A**). At day 4 post-treatment, we harvested tumors and measured viral antigen by immunofluorescence. Viral antigen was highly co-localized with F4/80+ cells in mice treated with r3LCMV, suggesting that macrophages were preferentially infected with r3LCMV (**Fig. 1B**). We also interrogated whether r3LCMV could replicate in B16 melanoma cells. To answer this question, we incubated B16 melanoma cells for 48 hr with r3LCMV vectors expressing a reporter protein, green fluorescent protein (GFP) at an MOI of 0.1, and then we measured viral antigen at day 2 by immunofluorescence. The r3LCMV vector was able to infect B16 melanoma cells in vitro, consistent with a prior study (*11*) (**Fig. S1**).

**Figure 1.**
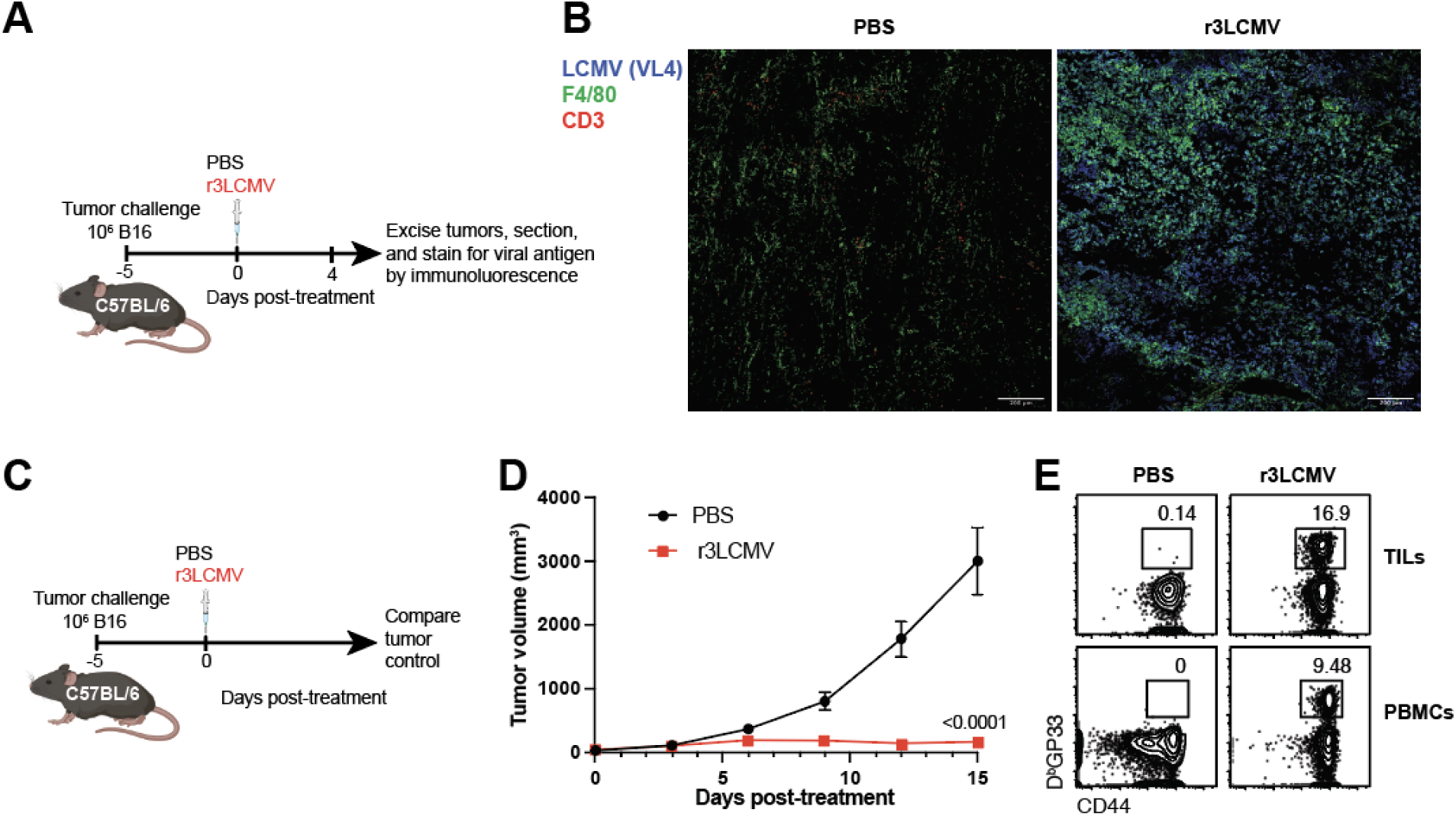
Attenuated r3LCMV replicates in B16 tumors and improves tumor control. (**A**) Experiment outline for evaluating whether r3LCMV replicates in tumor cells. (**B**) Representative immunofluorescence staining in tumor sections at day 4 post-treatment. (**C**) Experiment outline for evaluating whether r3LCMV improves tumor control. (**D**) Tumor control. (**E**) Representative FACS plots showing LCMV-specific CD8 T cell responses at day 7 post-treatment (gated on live CD8 T cells). Mice were treated intratumorally with 2x10^5^ PFU of r3LCMV, five days after tumor challenge. Data are pooled from 2 experiments (one experiment with n=5 per group and another with n=7 per group). Error bar represents SEM. Indicated P values were calculated by the Mann– Whitney test.

We also evaluated whether intratumoral treatment with r3LCMV improves tumor control. Interestingly, treatment of tumor-bearing mice with r3LCMV induced a significant improvement in tumor control (**Fig. 1C-1D**). To determine whether the antimoral effect of this LCMV vector was dependent on the ability of the virus to replicate, we injected B16 melanoma-bearing mice with replicating or non-replicating LCMV vectors (both were attenuated relative to wild type LCMV). We utilized a bisegmented (rLCMV) vector that can enter cells and express viral proteins but cannot induce a second round of infection due to a genetic absence of the glycoprotein (GP) gene, the viral protein that mediates viral entry. During the *in vitro* production of this bisegmented rLCMV vector, the GP is only provided *in trans* in the producer cells to allow entry of the vector into host cells, but progeny virions are unable to form infectious progeny virions due to genetic lack of the GP. On the other hand, the r3LCMV vector expresses GP in cis, allowing it to undergo several replication cycles until it is eliminated by the host’s immune response , but it is still significantly attenuated and does not replicate to wild type levels (*12*). Interestingly, the replicating (r3LCMV) vector resulted in a superior antitumoral effect relative to non-replicating (rLCMV) vector (**Fig. 2A-2D**).

**Figure 2.**
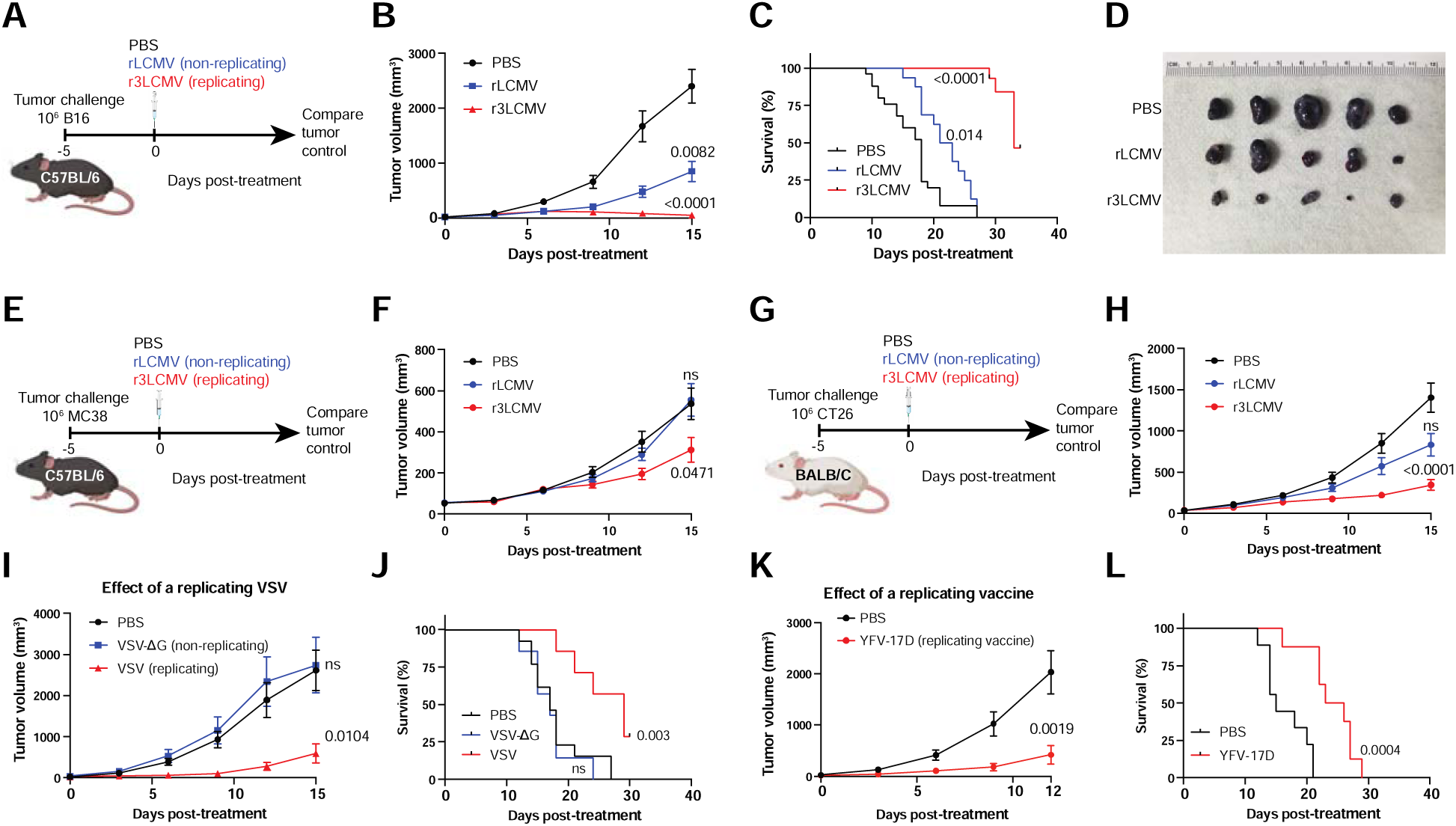
Viral replication is critical for the antitumoral effect of r3LCMV. (**A-D**) Effect of replicating r3LCMV versus non-replicating rLCMV vectors in the B16 melanoma model in C57BL/6 mice. (**A**) Experiment outline. (**B**) Tumor control. (**C**) Survival. (**D**) Representative images of tumors at day 8 post-treatment. (**E-F**) Effect of replicating versus non-replicating LCMV vectors in the MC38 colon carcinoma model in C57BL/6 mice. (**E**) Experiment outline. (**F**) Tumor control. (**G-H**) Effect of replicating versus non-replicating LCMV vectors in the CT26 melanoma model in BALB/c mice. (**G**) Experiment outline. (**H**) Tumor control. (**I-J**) Effect of replicating versus non-replicating VSV in the B16 melanoma model in C57BL/6 mice. (**I**) Tumor control. (**J**) Survival. (**K-L**) Effect of replicating YFV-17D vaccine in the B16 melanoma model in C57BL/6 mice. (**K**) Tumor control. (**L**) Survival. Mice were treated intratumorally with 2x10^5^ PFU of the indicated viruses, five days after tumor challenge. LCMV data are pooled from 2 experiments (one experiment with n=5 per group and another with n=10-12 per group). VSV data are pooled from 2 experiments (one experiment with n=3 per group, and another with n=4 per group). YFV-17D data are pooled from 2 experiments (one experiment with n=4 per group, and another with n=4-5 per group). Error bar represents SEM. Indicated P values were calculated by the Mann–Whitney test, or log rank test when comparing survival.

Intratumoral r3LCMV therapy also induced potent antitumoral effects in other tumor models, such as the MC38 colon carcinoma (**Fig. 2E-2F**), and in mice with different genetic backgrounds (**Fig. 2G-2H**), suggesting a generalizable antitumor effect. To a lesser extent, a replicating vesicular stomatitis virus (VSV), and the replicating yellow fever virus (YFV-17D) vaccine also induced antitumoral effects (**Fig. 2I-2L**), suggesting that the antitumoral effects of viral vectors are dependent of viral replication. We also tested antitumoral effects on distal tumors, also known as abscopal effect. To test this, we injected both flanks of the mice with B16 melanoma cells, followed by intratumoral r3LCMV injection in the right tumor only. Interestingly, r3LCMV also induced partial regression of the contralateral (left) tumor (**Fig. S2**), demonstrating the induction of abscopal effect. Further, we evaluated whether r3LCMV vectors synergized with PD-L1 blockade (**Fig. S3A**). Treatment with r3LCMV alone resulted in significantly superior antitumoral control relative to PD-L1 blockade alone, and there was a pattern of improved survival in mice that received combined treatment, relative to r3LCMV treatment alone, although the difference was not statistically significant (**Fig. S3B-S3C**).

In addition, we observed that treatment of tumor-bearing mice with r3LCMV induced a significant reduction of systemic and tumor-draining lymph node T regulatory cells (Tregs) after a week of treatment (**Fig. S4A-S4C**). Treg depletion has been shown to improve tumor control (*13*), so we examined whether Treg depletion could synergize with r3LCMV treatment. We utilized FoxP3-DTR mice, which allow for depletion of Tregs upon diphtheria toxin administration (*14*). Our data show that Treg depletion combined with r3LCMV treatment results in more potent antitumoral control, relative to Treg depletion alone (**Fig. S4D-S4E**).

### Role for antigen presentation, costimulation, and T cells

T cells are thought to be critical for the control of tumors, and their activation is dependent on two signals: antigen presentation and costimulation. We first examined the role of antigen presentation by challenging mice with β2-microglubulin KO (β2M-/-) B16 melanoma cells, which are unable to present antigen to CD8 T cells (*15*). β2M-/- B16 tumor-bearing mice were then treated with PBS or r3LCMV and tumor control was measured. Interestingly, β2M-/- B16 tumor-bearing mice treated with r3LCMV showed improved tumor control relative to control treated mice, suggesting that antigen presentation via MHC I was dispensable for the antitumoral effect (**Fig. 3A-3C**).

**Figure 3.**
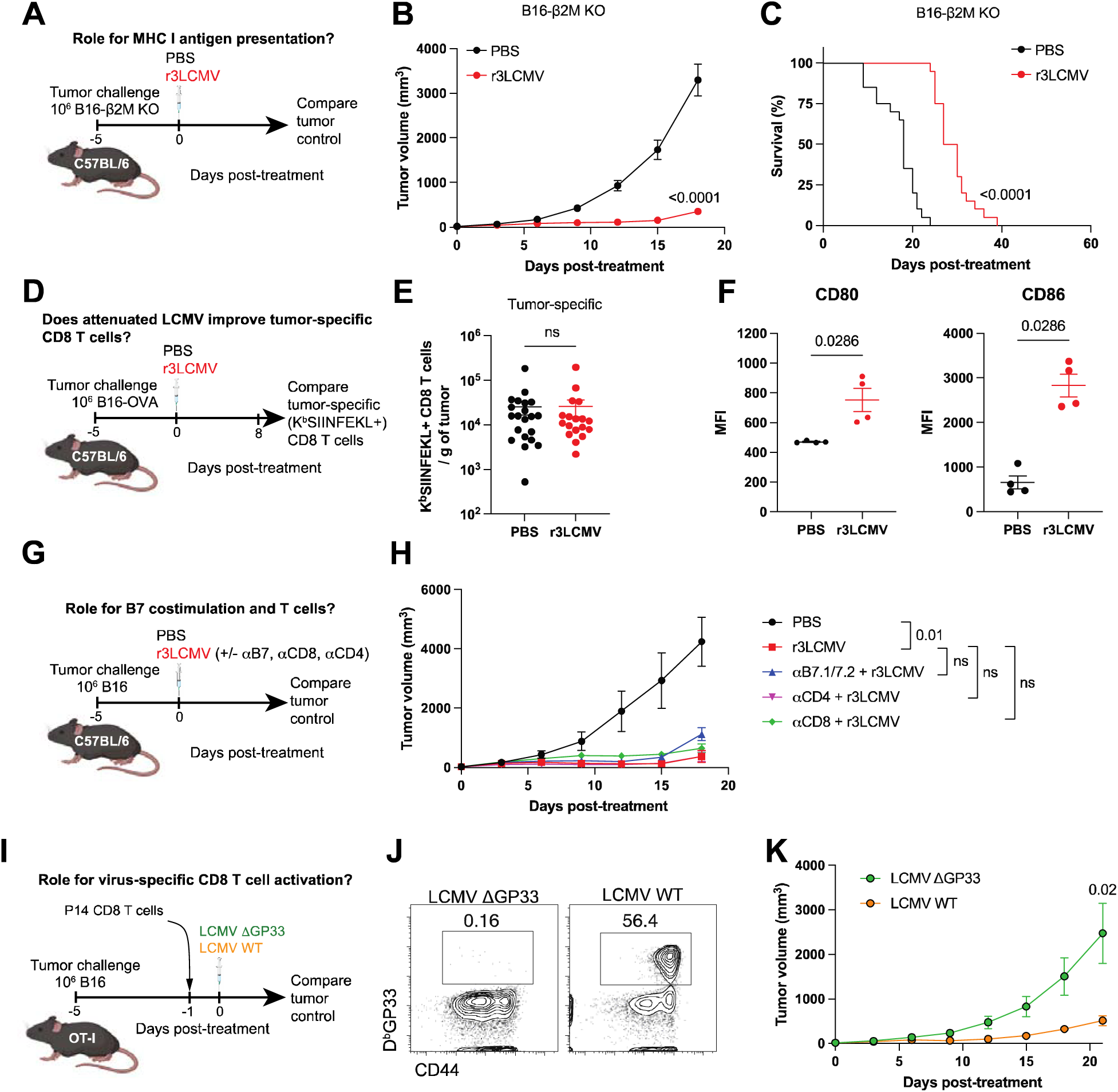
r3LCMV can exert antitumoral effects without the need for CD8 T cells and B7/CD28 costimulation. (**A-C**) Effect of r3LCMV vectors in the B16 β2M-/- melanoma model. (**A**) Experiment outline. (**B**) Tumor control. (**C**) Survival. (**D-E**) Effect of LCMV vectors on tumor-specific CD8 T cell responses. (**D**) Experiment outline. (**E**) Tumor-specific CD8 T cells at day 8 post-treatment. (**F-G**) Effect of r3LCMV vectors on B7 costimulatory molecules. (**F**) CD80 and CD86 costimulatory molecules on DCs from tumor-draining lymph nodes. DCs were gated on live, CD3-, NK1.1-, Ly6G-, CD19-, CD11b+, CD11c+ at day 4 post-treatment. (**G-H**) Effect of B7 costimulation blockade, CD8 T cell depletion, and CD4 T cell depletion. B7.1/B7.2 blocking antibodies, CD8 T cell depleting antibodies, or CD4 T cell depleting antibodies were administered intraperitoneally every 3 days. (**G**) Experiment outline. (**H**) Tumor control. (**I-K**) Effect of virus-specific CD8 T cells. (**I**) Experiment outline. (**J**) Representative FACS plots showing LCMV-specific CD8 T cell (P14) expansion in PBMCs at day 7 post-treatment. (**K**) Tumor control. Data from panels A-C are pooled from 2 experiments (one experiment with n=10 per group and another with n=10 per group). Data from panels D-E are pooled from 2 experiments (one experiment with n=9 per group and another with n=9-12 per group). Data from panel F are from 1 representative experiment (n=4 per group). Data from panels G-H are from 1 representative experiment (n=6-7 per group). Data from panels I-K are from 1 representative experiment (n=5-7 per group). Error bar represents SEM. Indicated P values were calculated by the Mann–Whitney test, or log rank test when comparing survival.

We then evaluated whether the r3LCMV treatment improves tumor-specific T cell responses, by challenging mice with B16 melanoma cells expressing OVA (B16-OVA), and then measuring OVA-specific CD8 T cell responses by tetramer staining. Although the r3LCMV treatment induced LCMV-specific CD8 T cell responses (**Fig. 1E**), it did not result in improvement of tumor-specific CD8 T cell responses (**Fig. 3D-3E**). We then measured costimulatory molecule expression on dendritic cells (DC) from tumor-draining lymph nodes following r3LCMV treatment, and we observed a significant increase in B7.1 (CD80) and B7.2 (CD86) molecule expression in mice that received r3LCMV treatment (**Fig. 3F**), suggesting a potential role for B7 costimulation. However, blockade of B7.1 and B7.2 molecules did not abrogate the antitumoral effect of r3LCMV (**Fig. 3G-3H**), suggesting that B7/CD28 costimulation was dispensable for the antitumoral effect of r3LCMV. We further examined the role for CD4 T cell responses by depleting these cells using depleting antibodies. CD4 T cell depletion did not impair the antitumoral effect of r3LCMV (**Fig. 3H**).

Moreover, we performed CD8 T cell depletion experiments to evaluate whether CD8 T cells were mechanistically involved. CD8 T cell depletion did not significantly impact the antitumoral effect of r3LCMV, but there was a pattern of reduced antitumoral control in mice that received CD8 T cell depleting antibodies and r3LCMV therapy, relative to r3LCMV therapy alone (**Fig. 3H**). This result motivated us to utilize an adoptive CD8 T cell transfer model to more rigorously measure the contribution of virus-specific CD8 T cells in tumor control. We transferred TCR transgenic CD8 T cells recognizing the LCMV GP33-41 epitope (P14 cells) into recipient tumor-bearing OT-I mice, which contain only OVA-specific CD8 T cells. This adoptive transfer model allowed us to examine the contribution of virus-specific CD8 T cell activation in a “T cell-replete” environment. We used OT-I mice as recipients instead of Rag1-/- mice because transferring donor T cells into Rag1-/- mice would lead to rapid homeostatic proliferation of donor T cells (emptiness-induced proliferation) (*16*). One day after P14 cell transfer, recipient mice were infected intratumorally with an LCMV variant lacking the GP33-41 epitope (LCMV ΔGP33) or a wild-type “parental” LCMV (Cl-13) to determine whether the activation of virus-specific CD8 T cells potentiates tumor control (**Fig. 3I**). Both LCMV strains replicate at similar levels and their only difference is a V→A escape mutation that destroys the GP33 epitope recognized by P14 cells (*17*). As expected, intratumoral infection with the wild type LCMV (but not the LCMV ΔGP33 variant) triggered robust P14 cell expansion in the tumor-bearing OT-I mice (**Fig. 3J**). Interestingly, the mice that were infected with the wild type LCMV (which showed robust P14 expansion) exhibited superior tumor control and survival relative to the mice that were infected with the LCMV ΔGP33 variant **(Fig. 3K**), suggesting that “bystander” activation of virus-specific CD8 T cells can facilitate tumor control in a host devoid of tumor-specific T cell responses. Collectively, these data using transgenic P14 cells suggested that the bystander activation of virus-specific CD8 T cells could potentiate tumor control.

### Intratumoral treatment with r3LCMV improves tumor control in the absence of adaptive immunity

To interrogate the role of adaptive immunity more rigorously, we challenged Rag1-/- mice with B16 tumor cells, followed by treatment with r3LCMV (**Fig. 4A**). Rag1-/- mice are unable to generate T cells and B cells (**Fig. 4B-4C**), leading to a severe combined immunodeficiency. Interestingly, Rag1-/- mice also exhibited a significant improvement in tumor control after r3LCMV treatment, demonstrating that r3LCMV could also induce antitumoral effects in the absence of adaptive immunity (**Fig. 4D**). We also examined whether the antitumoral effect of r3LCMV was dependent on IFNγ, also known as “adaptive interferon.” IFNγ is expressed mostly by effector T cells and this cytokine is important for tumor control (*18*). To examine the role of IFNγ on tumor cells, we challenged C57BL/6 mice with B16 melanoma cells lacking the IFNγ receptor (B16- IFNγR-/-). We then treated mice with PBS or r3LCMV to examine whether tumor control by r3LCMV therapy was dependent on tumor-intrinsic IFNγ signaling. Importantly, tumor control by r3LCMV was not dependent on IFNγ sensing by the tumor (**Fig. S5**), suggesting that tumor control by r3LCMV therapy was not dependent on “adaptive” interferon.

**Figure 4.**
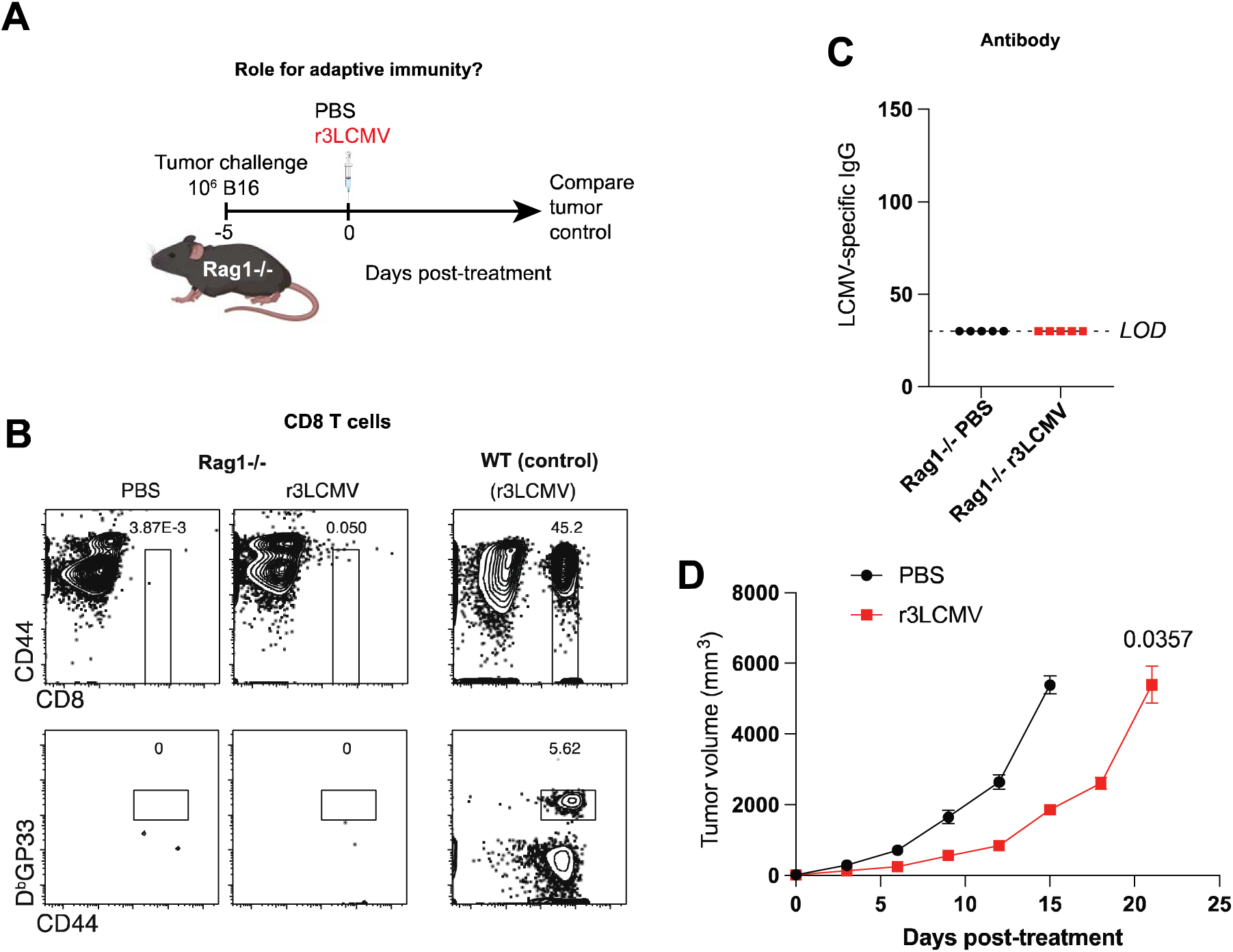
r3LCMV therapy improves tumor control in Rag1-/- mice. (**A**) Experiment outline. (**B**) Representative FACS plots showing absence of CD8 T cell responses in Rag1-/- mice. (**C**) ELISA plots showing absence of antibody responses in Rag1-/- mice. (**D**) Tumor control. Data are from 1 experiment (with n=5 per group). Error bar represents SEM. Indicated P values were calculated by the Mann–Whitney test.

**Figure 5.**
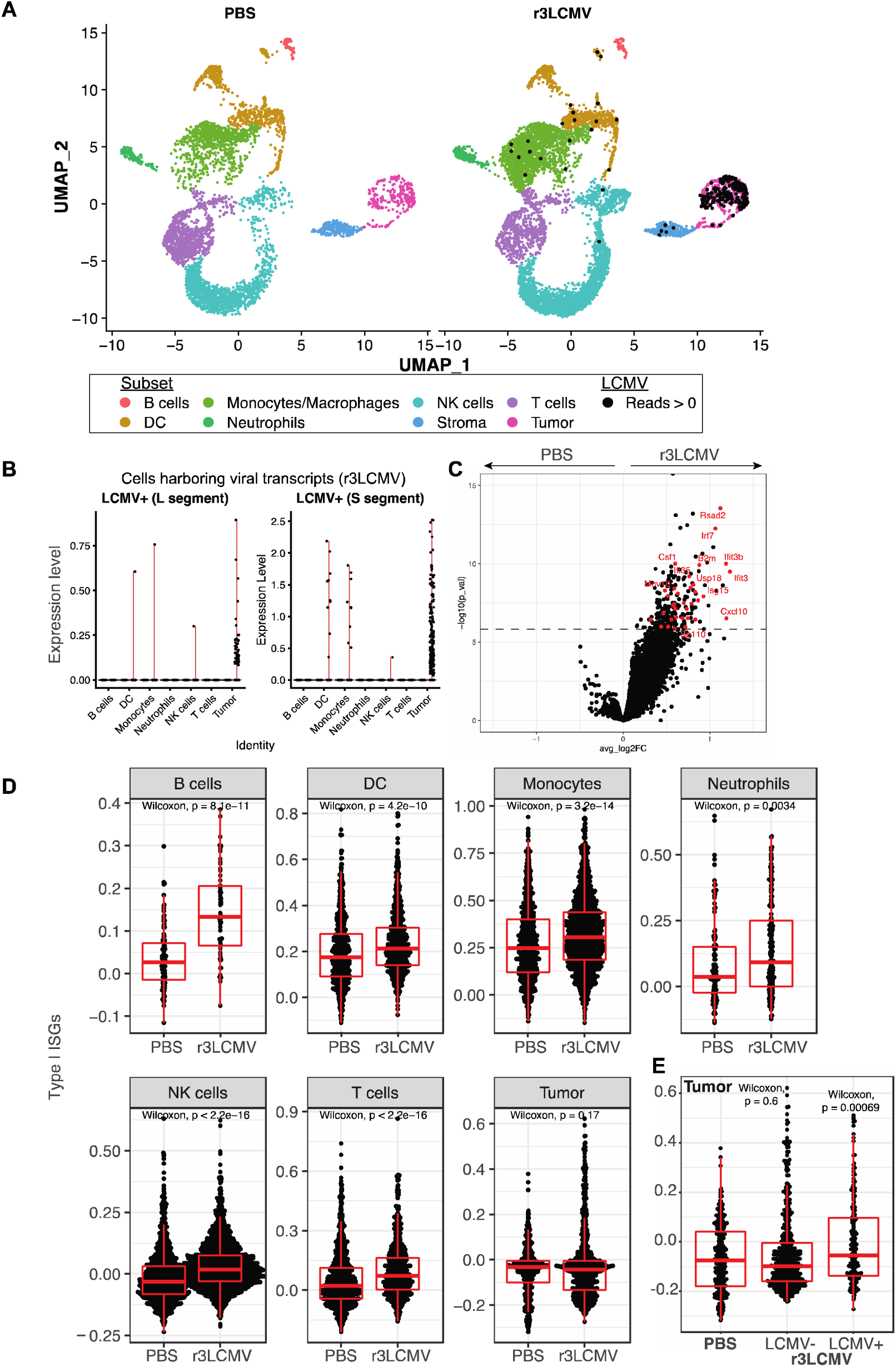
scRNA-Seq reveals enrichment of IFN-I responses by r3LCMV therapy. (**A**) UMAP plots showing cell distribution based on RNA expression. Each cell is colored by its inferred subset (based on ImmGen database). Cells harboring LCMV reads are indicated by a black dot. (**B**) Level of expression of LCMV L and S transcripts on different cell subsets from r3LCMV treated mice. (**C**) Volcano plot showing the differential expression of genes in tumor cells harboring LCMV reads versus those without LCMV reads. The dash line indicates p-value adjusted for multiple testing of 0.05. ISG are indicated in red. (**D**) Enrichment for ISG in different cell subsets. (**E**) ISG on tumor cells harboring LCMV or not harboring LCMV. This panel shows that tumor cells with LCMV reads express higher levels of ISG relative to tumor cells without LCMV reads. For each boxplot, the vertical line indicates the median, the box indicates the interquartile range, and the whiskers indicate 1.5 times the interquartile range. ∼80% of cells were CD45+ (after MACS purification). Each group represents pooled tumors from 5 different mice. Indicated P values were calculated by the Wilcoxon test.

### Single cell RNA-Seq (scRNA-Seq) analyses reveal a role for type I interferons

We then performed gene expression analyses to understand the effects of r3LCMV on different cell subsets within the tumor microenvironment. We harvested tumors at day 4 post-treatment, followed by scRNA-Seq analyses. We observed substantial differences in cell populations between the PBS and r3LCMV treated mice. Our single cell gene expression data show that r3LCMV induces changes in cell frequencies within the tumor microenvironment, including a significant influx of NK cells and macrophages (**Fig. 5A**). LCMV viral reads were detected in r3LCMV treated mice, especially in DCs, macrophages, and tumor cells themselves (which harbored the L and S RNA segments from LCMV) (**Fig. 5A-5B**). These gene expression data also show that r3LCMV induces several IFN-induced genes (ISG), including those coding for master transcription Irf7 and the antiviral protein Ifi3 and Isg15 (**Fig. 5C**). ISG were significantly upregulated in immune cell subsets; but not in tumor cells when analyzed as a whole (when compounding uninfected and infected tumor cells) (**Fig. 5D**). However, when we analyzed tumor cells containing viral transcripts versus tumor cells lacking viral transcripts in the r3LCMV treated mice, we observed significant upregulation of ISG only in tumor cells containing viral transcripts (**Fig. 5E**). These data suggested a possible role for IFN-I in the antitumoral control elicited by r3LCMV.

We then validated the gene expression data at the protein level. IFN-I and interferon-induced cytokines were highly upregulated in the plasma of mice treated with r3LCMV (**Fig. 6A**), consistent with other studies examining cytokine responses with other LCMV vectors (*7, 12*). We also performed mechanistic validation of our scRNA-Seq data. In particular, we evaluated the mechanistic roles of IFN-I by treating tumor-bearing mice with an IFN-I receptor–blocking antagonist (αIFNAR1 antibody, clone MAR1-5A3), which has been used in prior studies to block the IFN-I pathway (*12, 19–22*). Blockade of the IFN-I pathway significantly blunted the antitumoral efficacy of r3LCMV therapy (**Fig. S6**), suggesting that IFN-I could play a role in the antitumoral effect.

**Figure 6.**
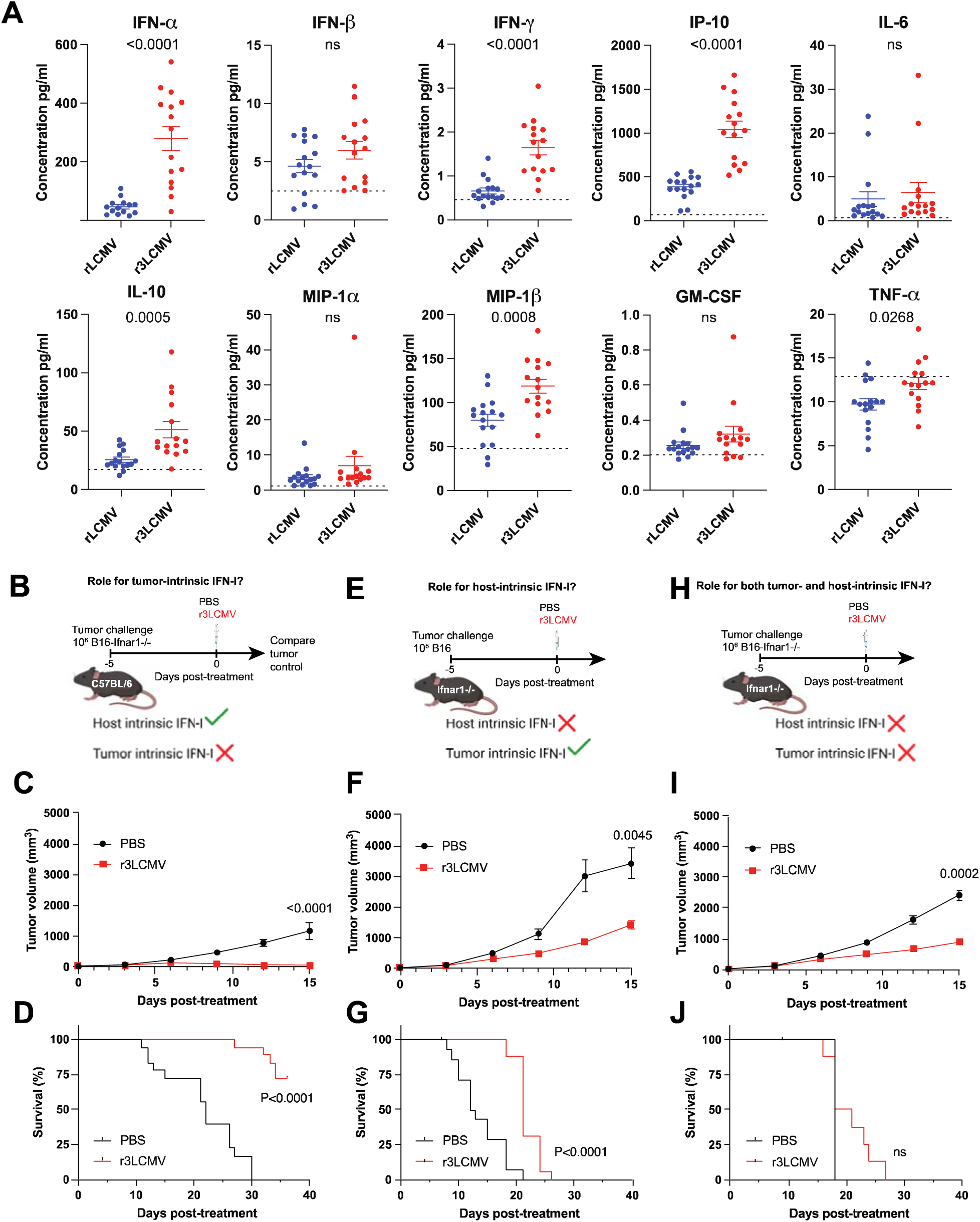
Confirmatory mechanistic studies corroborate a role for IFN-I. (**A**) Cytokine responses at day 1 post-treatment. Dashed lines represent naïve levels. (**B-D**) Effect of r3LCMV vectors on B16 IFNAR1-/- melanoma. (**B**) Experiment outline. (**C**) Tumor control. (**D**) Survival. (**E-G**) Effect of r3LCMV vectors on IFNAR1-/- mice. (**E**) Experiment outline. (**F**) Tumor control. (**G**) Survival. (**H-J**) Effect of r3LCMV vectors on B16 IFNAR1-/- melanoma in IFNAR1-/- mice. (**H**) Experiment outline. (**I**) Tumor control. (**J**) Survival. Data from panel A are pooled from 3 experiments (one experiment with n=5 per group, another with n=5 per group, and another with n=4-5 per group). Data from panels B-J are pooled from 2 experiments (one experiment with n=8-9 per group, and another with n=9-10 per group). Error bar represents SEM. Indicated P values were calculated by the Mann–Whitney test, or log rank test when comparing survival.

We performed three series of experiments to determine the tumor-intrinsic versus host-intrinsic roles of IFN-I. In our first model, we challenged mice with B16 melanoma cells lacking IFNAR1 (B16 *Ifnar1*-/-) (**Fig. 6B)**. These mice lacking IFN-I signaling specifically on tumor cells exhibited potent antitumoral responses and improved survival after r3LCMV treatment, suggesting that tumor-intrinsic IFN-I was dispensable (**Fig. 6C-6D)**. In our second model, we challenged Ifnar1-/- mice with B16 melanoma cells. In this model, where the host cells could not sense IFN-I, we observed that the antitumoral effect was modest and all mice succumbed within 4 weeks, suggesting that host-intrinsic IFN-I was important (**Fig. 6E-6G**). In our third model, we challenged Ifnar1-/- mice with B16 *Ifnar1*-/-melanoma cells. In this third model, where both the host and the tumor are unable to sense IFN-I, the antitumoral effect of r3LCMV was substantially dampened and there was no improvement in survival following r3LCMV therapy (**Fig. 6H-6J**). These data suggest that host-intrinsic IFN-I signaling is especially important for the antitumoral effect of r3LCMV therapy.

### r3LCMV treatment induces antitumoral effects without the need for NK cells and macrophages

Since the scRNA-Seq studies showed enrichment in NK cells and macrophages in the tumor upon r3LCMV treatment, we examined the contribution of these cells in tumor control. We first challenged mice with B16 melanoma tumors and then depleted NK cells continuously with an NK cell-depleting antibody to determine if this abrogated the antitumoral effect of r3LCMV. NK cell depletion did not abrogate tumor control by r3LCMV (**Fig. S7**). Similarly, continuous depletion of macrophages using clodronate liposomes did not abrogate tumor control after r3LCMV treatment (**Fig. S8**). Altogether, the antitumoral effect of r3LCMV did not appear to be dependent on NK cells and macrophages.

### r3LCMV induces a long-lasting antitumoral state

Tumors can recur throughout the lifespan of the host. In our experiments, all control PBS treated mice succumbed to the B16 melanoma challenge within weeks of challenge, whereas a fraction of r3LCMV treated mice typically survived. We interrogated whether mice that had cleared tumors (following r3LCMV therapy) developed immune memory to the tumor (following a secondary tumor challenge). Mice that were previously immunized with r3LCMV and that had cleared B16 tumors showed enhanced tumor control following a secondary tumor challenge, relative to control mice (**Fig. S9**), suggesting that r3LCMV induced a memory response to the tumor.

## Discussion

Therapeutic tumor vaccines based on recombinant viruses expressing a tumor antigen payload and oncolytic viruses that lyse tumor cells have emerged as promising anticancer agents due to their ability to activate innate and adaptive immune responses. For example, enteric cytopathic human orphan virus type 7 (ECHO-7) is currently being used for melanoma due to its ability to lyse tumor cells (*23*). In addition, an adenovirus-based vector is used for head and neck cancer; and an HSV-based vector is used for recurrent melanoma (*2, 24*). Less work has been done with nonlytic viruses that do not express any tumor antigen payload. Recently, LCMV vectors encoding HPV16 antigens started clinical testing for the treatment of HPV-related cancers. LCMV is a non-lytic arenavirus in clinical development as a vaccine vector to deliver tumor antigens to the immune system. LCMV does not directly lyse tumor cells, but it induces potent innate and adaptive immune responses. LCMV is also relatively proficient at evading antibody responses, allowing its re-utilization as a vaccine vector in a seropositive host (*8, 25*).

Prior research has shown that LCMV vectors can outperform protective efficacy elicited by other viral vector platforms, including Ad5 and poxvirus vectors (*7, 26*). When tumor-bearing mice are immunized with LCMV vectors containing the tumor antigen, they demonstrate stronger antitumor control relative to mice immunized with adenovirus or poxvirus vectors containing the same tumor antigen. However, ongoing clinical trials with LCMV vectors engineered to express tumor antigens are not assessing the potential contribution of tumor non-specific responses or whether immune activation by the vector itself can modulate tumor control.

Prior studies using models of therapeutic vaccination have shown that tumor-specific T cells play a critical role in LCMV vectored cancer therapy (*7, 11*), but it remains uncertain whether LCMV vectors that do not express any tumor antigen can also exert antitumoral activity. Historically, the use of the viral vectors as cancer vaccines requires knowledge of the specific neoantigens encoded by the tumor, which may vary between different patients and tumor types. In our study, we utilized a “generic” r3LCMV platform that does not encode any tumor antigen. Since the r3LCMV vector and the tumor do not share any antigen, the antitumoral effect that we report is considered bystander. Earlier studies by Lang and others have shown that infection of tumor-bearing mice with chronic strains of LCMV can improve tumor control (*27–30*). However, safety concerns of using live LCMV have deemed this approach not translatable to humans. A prior clinical trial used live LCMV in cancer patients, but these patients died with evidence of multi-organ LCMV infection upon necropsy (*31*). However, these patients were in the late stages of lymphoma and it is unclear if for that reason they succumbed; one patient harbored bacterial infection at the time of death and it remains unknown if death was caused by LCMV itself. Chronic LCMV infection can also render the host more susceptible to other diseases due to immune exhaustion caused by infection (*32*). Therefore, attenuated replicating LCMV vectors represent a safer clinical approach, given their high immunogenicity despite their limited ability to replicate. We report their safety and efficacy even in Rag1-/- mice.

In our study, we used an attenuated r3LCMV vector that replicates substantially lower than the parental virus, but is still able to trigger a robust innate and adaptive immune response. Very low levels of systemic virus can be detected 72 hr after infection with attenuated r3LCMV, with mice showing only a very transient viremia near the limit of detection (<5 PFU/mL) with no weight loss or signs of disease (*12*). Our studies show that T cells, B cells, NK cells, and macrophages (as well as other phagocytes that can be depleted by clodronate liposomes like monocytes, dendritic cells, and neutrophils (*33*)) are not necessary for the antitumoral effect of r3LCMV. However, we found that activation of virus-specific CD8 T cells can facilitate the control of the tumors, as shown by our P14 adoptive transfer studies. In summary, we demonstrate that attenuated r3LCMV vectors exert antitumoral effects partly via host-intrinsic IFN-I and that they are safe and effective even in an immunodeficient host. These studies are important for the development of LCMV-based therapies for cancer and for improving the mechanistic understanding of how nonlytic viral vectors modulate tumor immunity.

## Limitations of the study

Absence of host-intrinsic IFN-I signaling limits the antitumor efficacy of r3LCMV, yet mice devoid of host-intrinsic IFN-I signaling show partial tumor control upon r3LCMV treatment, suggesting that other immune pathways may contribute to the antitumoral effects. Future studies will examine the contribution of other innate immune pathways in the antitumoral effects elicited by r3LCMV.

## Acknowledgements

We thank Dr. Juan Carlos De La Torre (Scripps Research Institute, La Jolla, CA), Dr. Stephen Waggoner and Dr. Carolyn Rydyznski (Cincinnati Children’s Medical Center, Cincinnati, OH) for technical assistance making the r3LCMV vectors. We thank Drs. Jennifer Wu, Bin Zhang, and Chyung-Ru Wang for discussions. We also thank Hookipa Biotech for sharing rLCMV vectors and for helpful discussions. We thank Dr. Rebecca Obeng for guidance on the tissue sectioning and immunostaining of tumor samples. Several images were created with Biorender.com. This work was possible with a grant from the National Institute on Drug Abuse (NIDA, DP2DA051912) to P.P.M., an Institutional Research Grant (IRG-15-173-21) from the American Cancer Society to P.P.M., a pilot grant from the Lurie Cancer Center at Northwestern University.

## Author Contributions

Y.R.C and B.A. performed most of the mouse tumor challenge and immunogenicity experiments. T.D. helped to perform some of the mouse experiments. S. F. analyzed the gene expression data. P.P.M. designed the experiments and secured funding.

P.P.M. wrote the paper with feedback from all authors. The gene expression analyses in this research was supported in part through the computational resources and staff contributions provided by the Genomics Compute Cluster which is jointly supported by the Feinberg School of Medicine, the Center for Genetic Medicine, the Feinberg’s Department of Biochemistry and Molecular Genetics, the Office of the Provost, the Office for Research, and the Northwestern Information Technology.

## Supplemental Figure Legends

**Figure S1.**
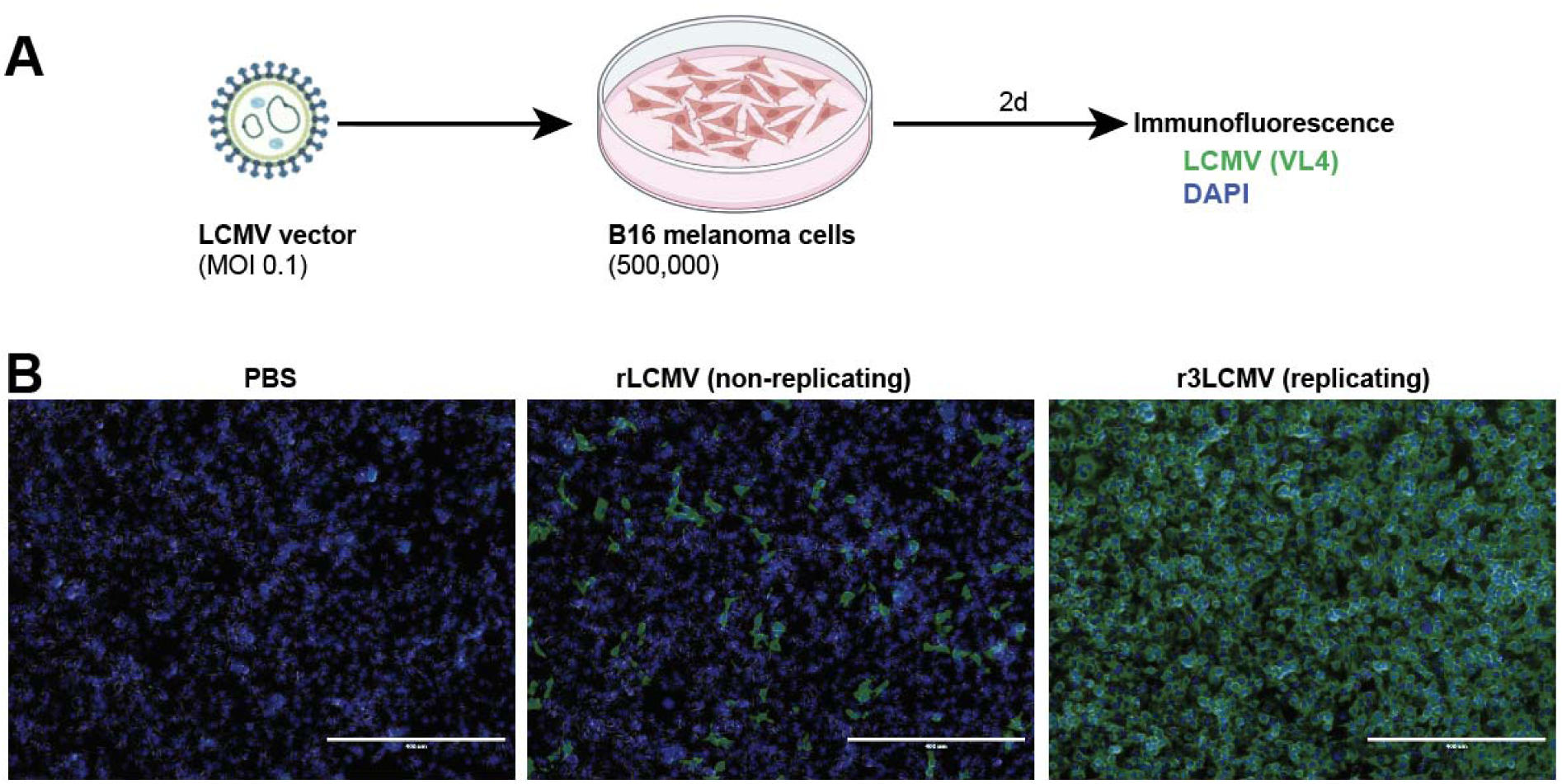
Attenuated r3LCMV replicates in B16 melanoma cells. (**A**) Experiment outline for detecting viral antigen after in vitro infection of B16 melanoma cells with replicating (r3LCMV) or non-replicating (rLCMV) vectors. (**B**) Representative immunofluorescence staining in B16 monolayers at day 4 post-infection. In this experiment, we detected substantially more viral antigen in B16 monolayers that were infected with replicating (r3LCMV), relative to non-replicating (rLCMV) vector. Experiment was performed 2 times with similar results.

**Figure S2.**
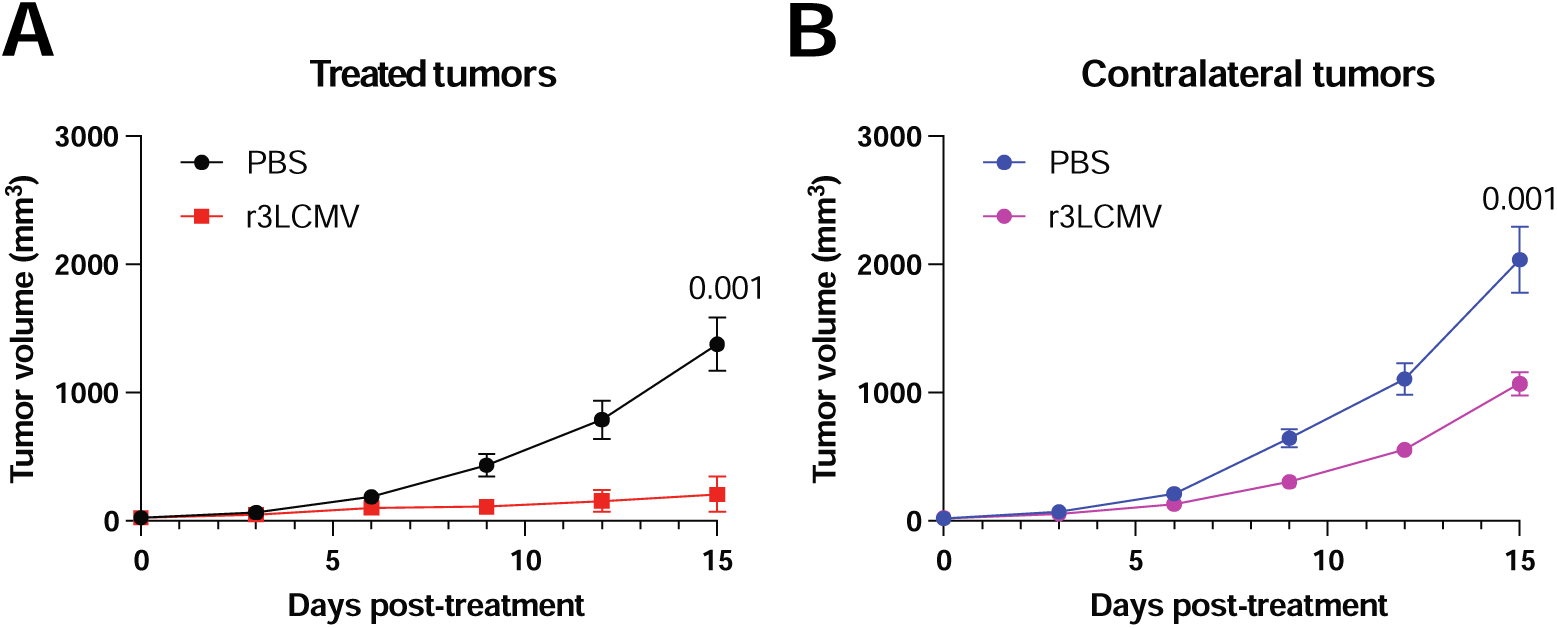
Attenuated r3LCMV induces abscopal effect. Mice were treated intratumorally with 2x10^5^ PFU of r3LCMV on the left tumor, five days after bilateral tumor challenge. (**A**) Tumor control on the treated tumor (left side). (**B**) Tumor control on the contralateral untreated tumor (right side). Data are from 2 experiments with a total of n=9-10 per group. Error bar represents SEM. Indicated P values were calculated by the Mann–Whitney test.

**Figure S3.**
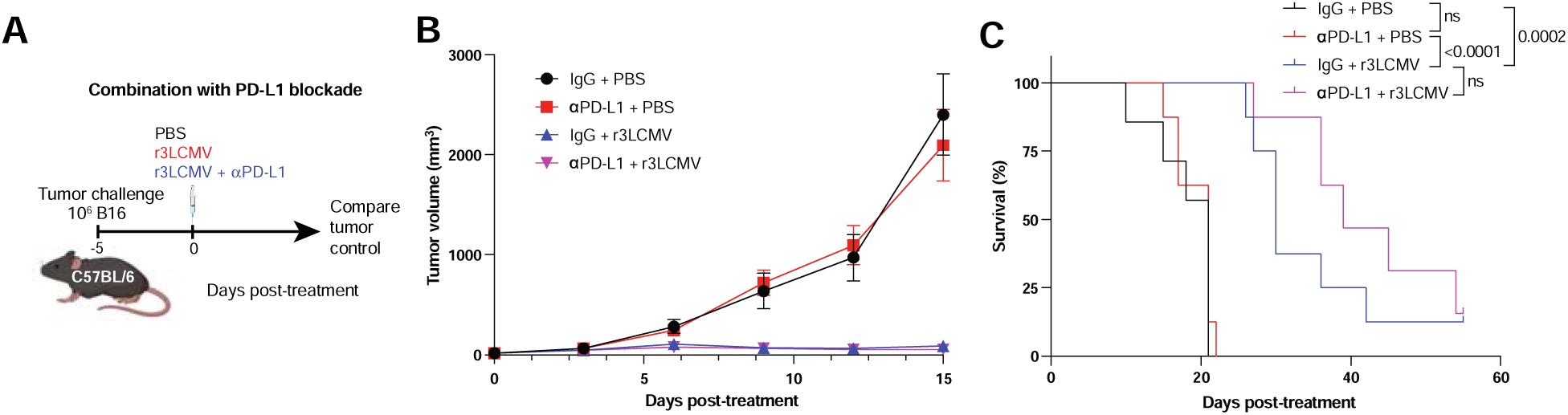
Effect of combined PD-L1 blockade and r3LCMV therapy. (**A**) Experiment outline for evaluating whether PD-L1 blockade improves r3LCMV therapy. (**B**)Tumor control. (**C**) Survival. Data are from 2 experiments with a total of n=6-7 per group. Error bar represents SEM. Indicated P values were calculated by the Kruskal-Wallis test and Dunn’s multiple comparison test, or Kaplan-Meier when comparing survival.

**Figure S4.**
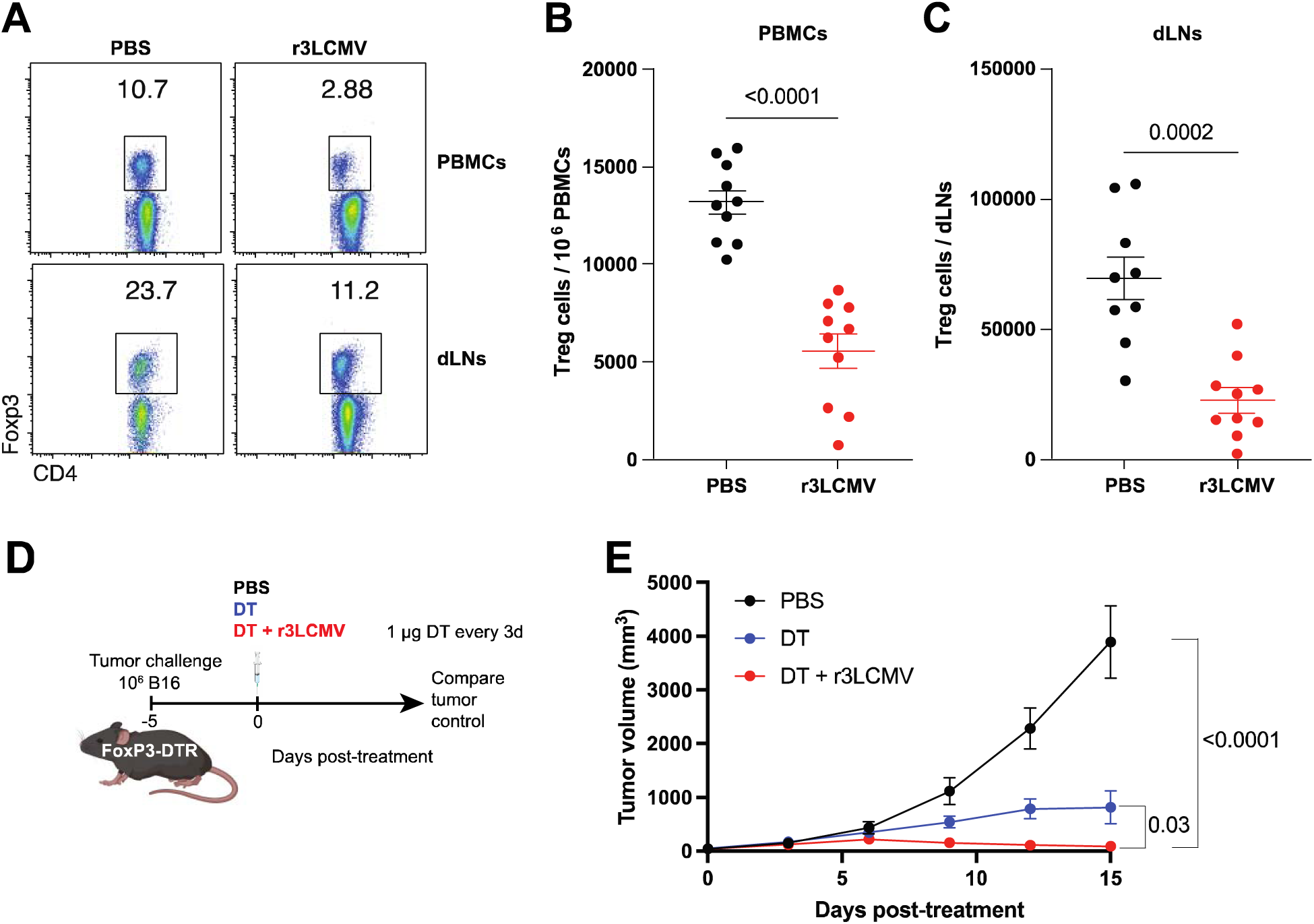
r3LCMV therapy results in a decline in Tregs. (**A**) Representative FACS plots showing Treg cell responses (gated on live CD4 T cells). (**B**) Summary of Treg cell responses in PBMCs. (**C**) Summary of Treg cell responses in tumor draining lymph nodes. Data from PBMCs are from day 7 post-treatment, and data from tumor draining lymph nodes are from day 8 post-treatment. (**D**) FoxP3-DTR mice were challenged with B16 melanoma tumors, similar to Fig. 1. After 5 days post-challenge, they were treated with diphtheria toxin (DT), with or without r3LCMV. (**E**) Tumor control. Data are pooled from 2 experiments (one experiment with n=5 per group and another with n=4-5 per group). Error bar represents SEM. Indicated P values were calculated by the Mann– Whitney test.

**Figure S5.**
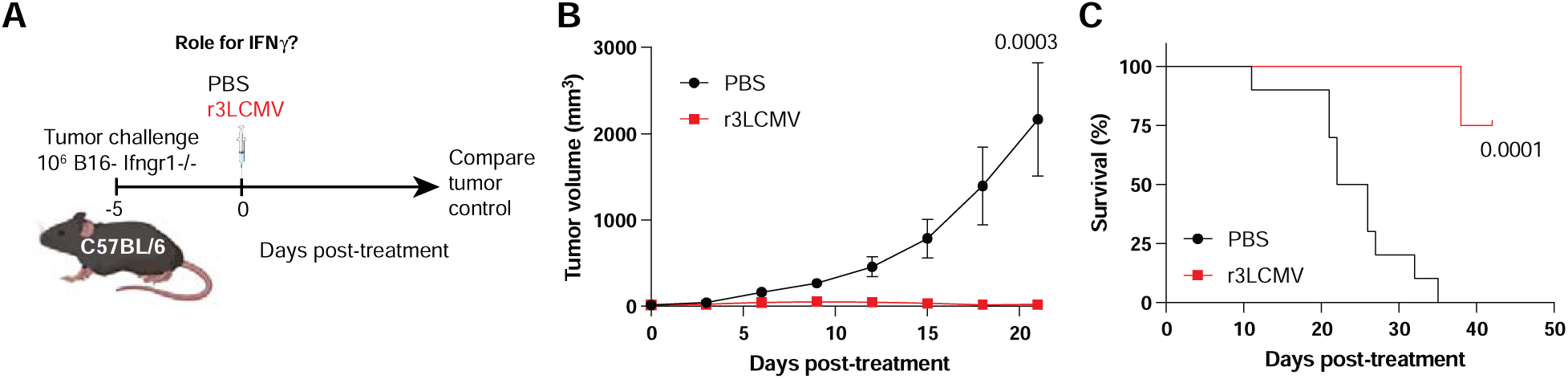
Tumor-intrinsic IFNγ signaling is not required for the antitumoral effect of r3LCMV. We tested the effect of r3LCMV vectors on B16 Ifngr1-/- melanoma. This tumor cannot sense IFNγ due to lack of its receptor. (**A**) Experiment outline. (**B**) Tumor control. (**C**) Survival. Data are from 1 experiment (n=8-9 per group). Error bar represents SEM. Indicated P values were calculated by the Mann–Whitney test, or Kaplan-Meier when comparing survival.

**Figure S6.**
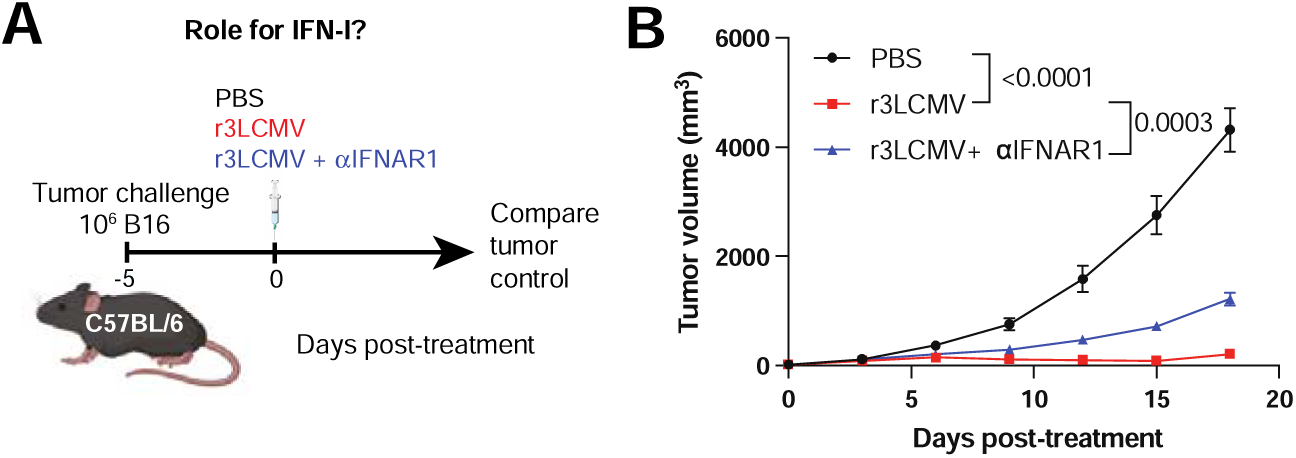
IFN-I signaling is partially required for the antitumoral effect of r3LCMV. We tested the effect of IFNAR1 blockade on r3LCMV therapy. (**A**) Experiment outline. (**B**) Tumor control. Data are pooled from 3 experiments (one experiment with n=6-7 per group, another with n=5, and and another with n=9-10 per group). Error bar represents SEM. Indicated P values were calculated using the Kruskal-Wallis test and Dunn’s multiple comparisons test.

**Figure S7.**
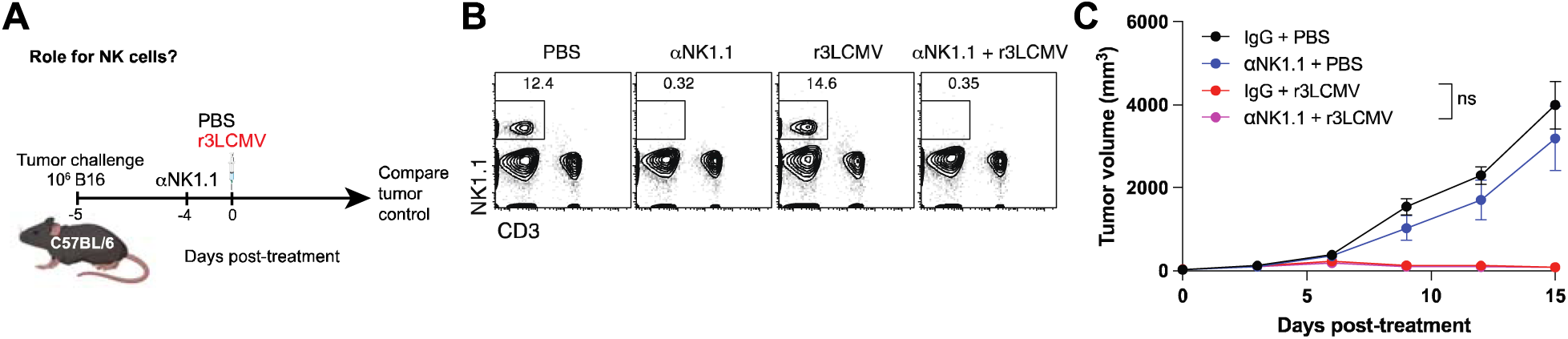
NK cells are not required for the antitumoral effect of r3LCMV. We tested the effect of NK cell depletion on r3LCMV therapy. (**A**) Experiment outline. (**B**) Representative FACS plots showing NK cells in PBMCs at day 0 of r3LCMV treatment (1 day after administration of αNK1.1). NK cell depleting antibodies (NK1.1, PK136) were administered at 500 μg, every 2 days, five times (see Materials and Methods). (**C**) Tumor control. Data are from 1 experiment (n=5 per group). Error bar represents SEM. Indicated P values were calculated using the Kruskal-Wallis test and Dunn’s multiple comparisons test.

**Figure S8.**
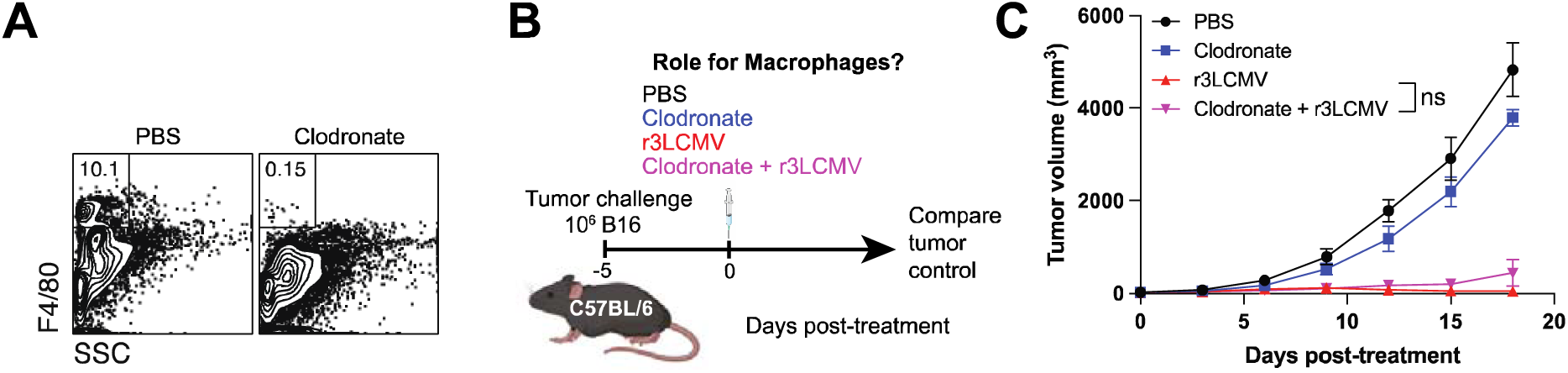
Macrophages are not required for the antitumoral effect of r3LCMV. We tested the effect of macrophage depletion on r3LCMV therapy. (**A**) Representative FACS plots of a pilot experiment showing macrophages in spleen at day 1 post-treatment (clodronate liposomes). This pilot showed that treatment with 200 μg of clodronate liposomes results in effective depletion of splenic macrophages. We used this same dose of clodronate liposomes. (**B**) Experiment outline. Clodronate liposomes were administered at 200 μg every 3 days, four times (see Materials and Methods). (**C**) Tumor control. Data are from 1 experiment (n=4-5 per group). Error bar represents SEM. Indicated P values were calculated using the Kruskal-Wallis test and Dunn’s multiple comparisons test.

**Figure S9.**
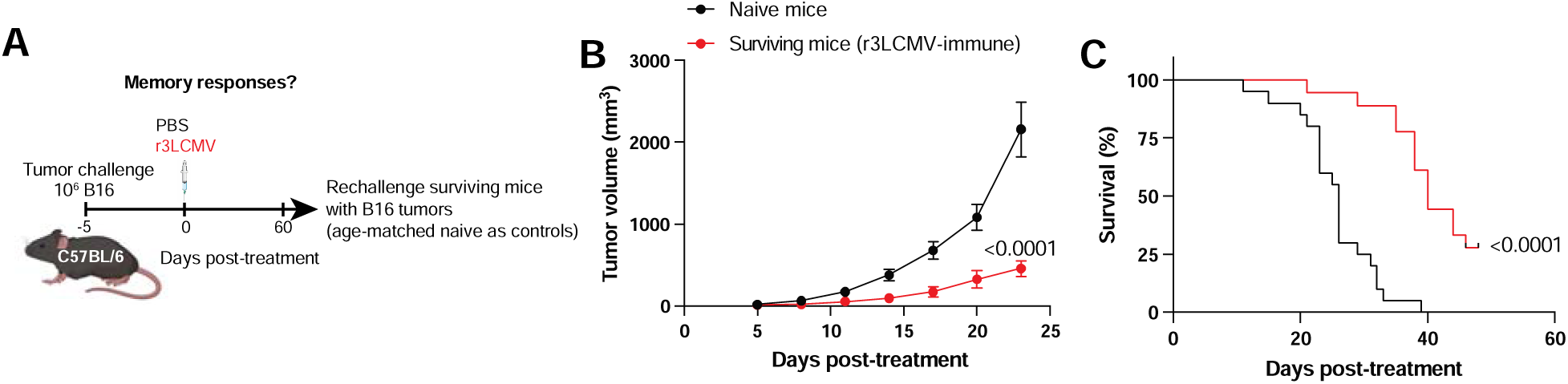
Immune memory after r3LCMV treatment. We tested whether mice that cleared tumors after r3LCMV therapy were protected upon subsequent tumor challenges. (**A**) Experiment outline. (**B**) Tumor control. (**C**) Survival. n=8-9 per group. Error bar represents SEM. Indicated P values were calculated using the Kruskal-Wallis test and Dunn’s multiple comparisons test.

## Materials and Methods

### Mice, tumor challenges, and LCMV vector treatments

Experiments were performed with 6-8-week-old wild type mice (half males and half females) from Jackson laboratories, Bar Harbor, ME (C57BL/6, Stock No: 000664; BALB/c, Stock No: 000651; IFNAR1-/-, Stock No: 028288; Rag1-/-, Stock 002216). Mice were challenged subcutaneously with 10^6^ B16 melanoma cells, MC38 colon carcinoma cells, and CT26 colon carcinoma cells and r3LCMV treatments started at day 5. Tumor volume was calculated as follows: Length x Width x Width x 1/2. Mouse challenges were performed at Northwestern University following BL2 guidelines with approval by the Institutional Animal Care and Use Committee (IACUC).

#### Cells and viruses

We used a murine melanoma cell line: B16 (gift from Dr. Chyung-Ru Wang at Northwestern University, Chicago); B16-OVA melanoma cell line (gift from Dr. Jennifer Wu at Northwestern University, Chicago); B16-β2M KO cells (gift from Dr. Omar Abdel-Wahab at Memorial Sloan Kettering Cancer Center, NY), B16-IFNAR1 KO and B16-IFNGR1 KO cells (Invivogen), MC38 (ATCC), and CT26 (ATCC). The tumor cells were cultured in DMEM (GIBCO, Cat# 11965-092) with 10% Fetal Bovine Serum (Sigma-Aldrich, Cat# F0926), 1% L-Glutamine (GIBCO, Cat# 25030-081), and 1% Penicillin and Streptomycin (GIBCO, Cat#15140-122) in 37°C 5% CO_2_ incubator. BHK-21 cells (ATCC, Cat# CCL-10) were used for production of LCMV, VSV, and YFV-17D. Vero E6 cells (ATCC, Cat# CRL-1586) were used for titration of r3LCMV, VSV, and YFV-17D. BHK-21 and Vero E6 cells were cultured in EMEM (ATCC, Cat# 30-2003) with 10% FBS, 1% L-Glutamine, and 1% Penn/Strep in 37 °C 5% CO_2_ incubator. Non-replicating (rLCMV) vectors expressing GFP (used in Figs. 2A-2H, 6A and Figure S1) were a kind gift from Hookipa Pharma Inc. For the rest of the experiments using attenuated vectors, we used replicating (r3LCMV) vectors expressing GFP, which were made using DNA plasmids from Dr. Juan Carlos De La Torre (Scripps Research Institute, La Jolla, CA). The LCMV strain lacking the GP33-41 epitope (GP35V→A escape mutation, which cannot be recognized by P14 cells) was derived from a prior study (*17*). This LCMV variant was used to examine the role of virus-specific CD8 T cell activation in tumor control.

### Adoptive cell transfer

CD8 T cells were purified from spleens of transgenic P14 mice, using a negative selection isolation kit (STEMCELL Technologies), and purity was confirmed to be >97%. 5x10^6^ CD8 T cells were injected into a mouse intravenously, one day before viral infection.

### Antibody treatments, cell depletions

All Antibodies for *in vivo* treatments were purchased from BioXCell or Leinco, and were diluted in sterile PBS and injected intraperitoneally (i.p.). PD-L1 blocking antibodies (10F.9G2) were administered at 200 μg, every three days, five times, as previously shown (*34*). B7.1 and B7.2 blocking antibodies (16-10A1 and GL-1, respectively) were administered at 200 μg each, every three days. IFNAR1 blocking antibodies (MAR1-5A3) were administered at 200 μg, every three days. This MAR1-5A3 antibody binds to interferon α/β receptor subunit 1 (IFNAR1) and blocks binding to interferons α/β, abrogating the induction of ISGs *in vivo* (*20, 22, 35–37*). NK cell depleting antibodies (NK1.1, PK136) were administered at 500 μg, every 2 days, five times. CD4 T cell depleting antibodies (GK1.5) and CD8 T cell depleting antibodies (2.43) were administered at 200 μg, every three days. Diphtheria toxin (Sigma-Aldrich) was administered at 1 μg i.p. (diluted in PBS), on days 0, 1, 4, 7, and 10 of r3LCMV therapy. This dose was similar to prior studies, using Foxp3-DTR knock-in mice on a C57BL/6 background (*14, 38*). Clodronate liposomes were administered at 200 μg every 3 days, four times.

### Quantification of viral titers

5x10^5^ of Vero E6 cells were plated onto each well in 6-well plates, and after 24∼48 hours when they reached ∼95% confluency, the media were removed and 200 μL of serial viral dilutions were added dropwise on top of the monolayer of the cells. Plates were rocked every 10 min in a 37 °C, 5% CO_2_ incubator for 1 hour. 200 μL of media was aspirated out, and the monolayers were gently overlaid with a 1:1 mixture of 2x 199 media (20% FBS, 2% Pen/Strep, 2% L-glutamine) and 1% agarose at 37°C. After 4 days, a second overlay was added, consisting of a 1:1 solution of 2x 199 media, 1% agarose, and 1:50 of neutral red. Overlay was removed on day 5 and plaques were counted using a conventional light microscope.

### Flow cytometry

MHC class I monomers (K^b^SIINFEKL or D^b^GP33,) were used for detecting virus-specific CD8 T cells, and were obtained from the NIH tetramer facility located at Emory University. MHC I monomers were tetramerized in-house. Single cell suspensions were stained with live/dead fixable dead cell stain (APC-Cy7, Invitrogen, cat# L34976A), anti-mouse CD8α (clone: 53–6.7, PerCP-Cy5.5, BD Pharmingen, cat # 551162; clone: 53– 6.7, FITC, BD Pharmingen, cat# 553031; clone: 53–6.7, APC, eBioscience, cat# 17-0081-83), anti-mouse CD4, (clone: RM4-5, PE-Cy7, eBioscience, cat# 25-0042-82; clone: RM4-5, Pacific Blue, eBioscience, cat# 57-0042-82), anti-mouse CD44 (clone: IM7, FITC, BD Pharmingen, cat# 553133; clone: IM7, Pacific Blue, Biolegend, cat# 103020), anti-mouse CD80 (clone: 16-10A1, FITC, BD Pharmingen, cat# 553768), anti-mouse CD86 (clone: GL1, PE, BD Pharmingen, cat# 561963), anti-IFNAR1 (clone: MAR1-5A3, PE, Biolegend, cat# 127312), anti-mouse CD11b (clone: M1/70, Alexa Fluor 700, Biolegend, cat# 101222), anti-mouse CD11c (clone: N418, PerCP Cy5.5, Biolegend, cat# 117328; clone: N418, PE-Cy7, Biolegend, cat# 117318), anti-mouse CD90.1 (Thy1.1) (clone: HIS51, eFluor 450, eBioscience, cat# 48-0900-82; clone: HIS51, PE-Cy7, eBioscience, cat# 25-0900-82), anti-mouse CD90.2 (Thy1.2) (clone: 53-2.1, APC, eBioscience, cat# 17-0902-82), anti-mouse CD45.1 (clone: A20, PE-Cy7, Biolegend, cat# 110730; clone: A20, FITC, BD Pharmingen, cat# 553775), anti-mouse CD45.2 (clone: 104, PE, Biolegend, cat# 109808; clone: 104, FITC, BD Pharmingen, cat# 553772), anti-mouse TCR Va2 (clone: B20.1, PE, BD Pharmingen, cat# 553289), anti-mouse CD279 (PD-1) (clone: RMP1-30, PE, Biolegend, cat# 109104; clone: RMP1-30, PE-Cy7, Biolegend, cat# 109110; clone: RMP1-30, FITC, eBioscience, cat# 11-9981-85), anti-mouse CD274 (B7-H1, PD-L1) (clone: 10F.9G2, PE, Biolegend, cat# 124308), anti-mouse Foxp3 (clone: FJK-16s, APC, eBioscience, cat# 17-5773-82), anti-mouse CD25 (clone: 3C7, PerCP-Cy5.5, Biolegend, cat# 101911), anti-mouse B220 (clone: RA3-6B2, PerCP-Cy5.5, Biolegend, cat# 103236), and anti-mouse CD3 (clone: 17A2, Pacific blue, Biolegend, cat# 100214; clone: 17A2, Biotin, Biolegend, cat# 100244), anti-mouse Ly-6G (clone: RB6-8c5, Biotin, eBioscience, cat# 13-5931-85), anti-mouse NK1.1 (clone: PK136, Biotin, eBioscience, cat# 13-5941-85; clone: PK136, PE, BD Pharmingen, cat# 553165), anti-mouse CD19 (clone: eBio1D3, Biotin, eBioscience, cat# 13-0193-82), SA-BV421 (Biolegend, cat# 405225), SA-APC (Invitrogen, cat# S868), and SA-PE (Biolegend, cat# 405204). Flow cytometry samples were acquired with a Becton Dickinson Canto II or an LSRII and analyzed using FlowJo v10 (Treestar).

### Tumor sectioning and immunofluorescence

Tumors were fixed in PLP fixative solution for 24 hours at 4°C. The tumor samples were washed with PBS and cryoprotected for 24 hours at 4°C in a sucrose/PBS dilution. The fixed tissue samples were frozen in OCT on dry ice. Once the samples were frozen, they were kept in -80°C freezer until sectioning. The tissue samples were sectioned using a cryomicrotome with 10 μm thickness. The frozen sections were washed with PBS 2 times for 5 minutes each time and rinsed in 0.05% PBS-T. The slides were incubated with the blocking solution (PBS + 1% BSA + 5% goat serum) for 10 minutes. The slides were stained with the primary and the secondary antibodies in the blocking solution for 2 hours and 1 hour, respectively. VL4 antibody (BioXCell) was used to detect LCMV antigen. After the primary and the secondary antibody staining, the slides were washed 2 times with PBS-T. The slides were washed with water and mounted with Vector AntiFade mounting medium. Slides were imaged at the Center for Advanced Microscopy (CAM) Cell Imaging Facility and Nikon Imaging Center at Northwestern University.

### Multiplex cytokine/chemokine assay

The mouse peripheral blood samples were collected in 1.5ml tubes 24 hours post-infection of LCMV. The blood samples were centrifugated at 15000 rpm in 4°C to separate the serum samples. The serum samples were collected and frozen in -80°C until its use. Multiplex cytokines/chemokines kit for IFNα, IFNβ, IFNγ, IL-6, IL-10, TNFα, IP-10, MIP-1α, MIP-1β, and GM-CSF was purchased from Mesoscale Diagnostics LLC.

### LCMV-specific ELISA

Binding antibody titers were quantified using ELISA as described previously (*12, 39, 40*), using LCMV GP as coating antigen. Briefly, 96-well, flat-bottom MaxiSorp plates (Thermo Fisher Scientific) were coated with 1 μg/mL of GP for 48 hours at 4°C. Plates were washed 3 times with wash buffer (PBS plus 0.05% Tween 20). Blocking was performed with blocking solution (200 μl PBS plus 0.05% Tween 20 plus 2% BSA) for 4 hr at room temperature. 6 μl of plasma samples were added to 144 μl of blocking solution in the first column of the plate, 3-fold serial dilutions were prepared for each sample, and plates were incubated for 1 hr at room temperature. Plates were washed 3 times with wash buffer. Goat anti-mouse IgG antibody tagged with streptavidin-HRP (Southern Biotech, 7105-05) was diluted 1:400 in blocking buffer and incubated for 1 hr at room temperature. After washing plates 3 times with wash buffer, 100 μl/well SureBlue Substrate (SeraCare) was added for 1 min. The reaction was stopped using 100 μl/well KPL TMB Stop Solution (SeraCare). Absorbance was measured at 450 nm using a Spectramax Plus 384 (Molecular Devices).

### Single-cell RNA sequencing (scRNA-seq)

5 different mice treated with r3LCMV and 5 different mice treated with vehicle (PBS) were enriched for CD45+ cells and pooled for single-cell sequencing, separately. Single-cell libraries were generated using 10x Genomics 3’ kits. Cell Ranger (version 6.1.2) was used to demultiplex raw base call files (BCL) to FASTQ files and align reads to the Mouse genome (Ensembl version GRCm39 version 110) supplemented with LCMV genome (GenBank accession NC_004291.1 and NC_004294.1). For counting, Cell Ranger was run with the option to include reads spanning intron regions of genes during counting; all remaining default options were used. Count matrices were further analyzed in R (version 4.6.2), Bioconductor (version 3.17) and the R package Seurat (version 4.3.0.1). The R package SingleR (version 2.2.0) with the ImmGen reference was used to annotate the subset of each cell. Differential expression was performed by fitting a negative binomial generalized linear model to gene expression and a likelihood-ratio test for statical testing. Benjamini-Hoshberg adjustment was used to correct for multiple testing and cutoff of 5% false-positive was considered significant. scRNA-seq was performed at the Northwestern University NUSeq core.

### Statistical analysis

Statistical analyses are indicated on the figure legends. Statistical significance was established at p ≤0.05. Data were analyzed using Prism (Graphpad).

